# Single Cell Phenotyping Reveals Heterogeneity among Haematopoietic Stem Cells Following Infection

**DOI:** 10.1101/080416

**Authors:** Adam L MacLean, Maia A Smith, Juliane Liepe, Aaron Sim, Reema Khorshed, Narges M Rashidi, Nico Scherf, Axel Krinner, Ingo Roeder, Cristina Lo Celso, Michael PH Stumpf

**Affiliations:** Department of Life Sciences, Imperial College London, London, UK; Institute for Medical Informatics and Biometry, Technische Universitat Dresden, Dresden, Germany; Current address: Department of Mathematics, University of California Irvine, USA.; Current address: Department of Molecular Oncology, BC Cancer Agency, and Graduate Bioinformatics Training Program, University of British Columbia, Vancouver BC, Canada.; Current address: Max Planck Institute for Biophysical Chemistry, Göttingen, Germany.; Current address: Max Planck Institute of Molecular Cell Biology and Genetics, Dresden, Germany.

## Abstract

The haematopoietic stem cell (HSC) niche provides essential micro-environmental cues for the production and maintenance of HSCs within the bone marrow. During inflammation, haematopoietic dynamics are perturbed, but it is not known whether changes to the HSC-niche interaction occur as a result. We visualise HSCs directly in vivo, enabling detailed analysis of the 3D niche dynamics and migration patterns in murine bone marrow following *Trichinella spiralis* infection. Spatial statistical analysis of these HSC trajectories reveals two distinct modes of HSC behaviour: (i) a pattern of revisiting previously explored space, and (ii) a pattern of exploring new space. Whereas HSCs from control donors predominantly follow pattern (i), those from infected mice adopt both strategies. Using detailed computational analyses of cell migration tracks and life-history theory, we show that the increased motility of HSCs following infection can, perhaps counterintuitively, enable mice to cope better in deteriorating HSC-niche micro-environments following infection.

**Author Summary:** Haematopoietic stem cells reside in the bone marrow where they are crucially maintained by an incompletely-determined set of niche factors. Recently it has been shown that chronic infection profoundly affects haematopoiesis by exhausting stem cell function, but these changes have not yet been resolved at the single cell level. Here we show that the stem cell–niche interactions triggered by infection are heterogeneous whereby cells exhibit different behavioural patterns: for some, movement is highly restricted, while others explore much larger regions of space over time. Overall, cells from infected mice display higher levels of persistence. This can be thought of as a search strategy: during infection the signals passed between stem cells and the niche may be blocked or inhibited. Resultantly, stem cells must choose to either ‘cling on’, or to leave in search of a better environment. The heterogeneity that these cells display has immediate consequences for translational therapies involving bone marrow transplant, and the effects that infection might have on these procedures.

## Introduction

Haematopoietic stem cell (HSC) function is essential for the development and maintenance of the haematopoietic system [1,2]. It relies on both cell-intrinsic and cell-extrinsic factors and their interplay at the sites of haematopoiesis [3,4]. At the sites where haematopoiesis takes place — in adult mammals this is typically the bone marrow, and on occasion the spleen — it is believed that there exist stem cell niches; this term refers to the anatomical and functional influence on HSCs from cell-extrinsic factors. HSC niches have been extensively studied, and an increasingly complex picture is emerging of the stromal and hematopoietic cell types able to influence HSC function, and the molecular cross-talk between HSCs and neighboring cells [5–10]. Nevertheless, there are a host of direct and indirect lines of evidence that point to the existence of dedicated environments that enable, and are indeed crucial for, HSCs to carry out their function.

Vasculature, endothelial and mesenchymal stromal cells are all constituent members of the stem cell niche, and interact with HSCs via signalling mediated by cytokines including CXCL12 (stromal cell-derived factor) and SCF (stem cell factor) [3,11–16]. HSCs have been found to locate near both the arterioles in the area of endosteum, and the smaller sinusoidal capillary vessels that pervade the inner bone marrow cavity [7,9,10,17,18]. Although the extent of influence of osteoblastic cells (residing near the endosteum) has been contested, they are likely either a constituent of the niche or support the establishment of niches [19]. Resolving this spatial complexity and the cellular dynamics, requires *in vivo* spatio-temporal data at the level of single HSCs, including ideally also genomic/transcriptomic analysis; but simultaneous *in vivo* phenotyping (through live imaging) and molecular characterisation of cells currently remains beyond our grasp. A number of relevant studies have helped to reveal homeostatic and non-homeostatic HSC–niche behaviour [20–22]. Spatial data that describe cell migration patterns enable the testing of hypotheses regarding the relative locations of different cell types, the modes of cell movement, and the factors that determine the behaviour of migrating cells [23,24].

Resolving HSC–niche dynamics requires spatio-temporal data not only for homeostatic conditions, but also under perturbations. Infection directly perturbs cells of the haematopoietic system through the immune response, and, in addition, has recently been shown to directly affect undifferentiated haematopoietic stem and progenitor cells [25–28]. Thus it provides a suitable model that compliments other HSC perturbations (such as cancer [29] or ageing [10,30]). There are a host of important reasons for studying the effects of infection on haematopoiesis: many infections, in particular those with high morbidity profoundly affect the constituency of the blood, and for several diseases, an ensuing weakness of the immune response can have the most pronounced medium and long-term consequences for the health of the infected individual. Also for acute, but especially for chronic infections, we lack detailed insights into the mechanisms by which the infection affects the haematopoietic system; though the fact that the HSCs, at the top of the haematopoietic cascade, can be affected is beyond doubt. *How* HSCs respond to a challenge by an infectious agent, and whether they do so uniformly and deterministically, is at best partially understood.

The nematode *Trichinella spiralis* causes trichinosis, a nonlethal infection in mice following ingestion [31]. Parasitic larvae produced in the host migrate to the circulatory system causing initial tissue damage and inflammation that provokes a response from regulatory cytokines following the acute disease phase [32]. This in turn affects haematopoiesis in terms of number and frequency of heaematopoietic progenitors, and increases the engraftment ability of HSCs despite their numbers and cell cycle profiles remaining unchanged [21]. This was linked to changes in the behavious of engrafting HSCs [21].

While the importance of single cell molecular profiling is being widely recognised, it is also important to understand the *phenotypic* variability — here of HSC migration behaviour — at the single cell level. Here we characterise and quantify the changes in HSC-niche interactions triggered by infection, at the single cell level. Based on the results of [21] — capturing cell movement via 3D tracking — we subject these imaging data to detailed statistical analysis of the observed movement patterns. We reconstruct the trajectories of HSCs through the bone marrow space and find significant differences between HSCs from control and *Trichinella*-infected mice. Strikingly, HSCs from infected mice show higher levels of sustained migration as well as more pronounced cell-to-cell variability in migration behaviour than HSCs from control mice. Taken together, these data show that infection leads to greater heterogeneity in the behaviour of HSCs: a subpopulation of HSCs are likely to leave their current niche and explore larger regions of bone marrow space. Within the framework of life history theory [33,34], we are able to give conditions under which such migration of HSCs may benefit an individual by corresponding to a simple bet-hedging strategy [35,36], which enables HSCs to move from deteriorating niches into other, more supportive niches. Thus phenotypic heterogeneity among a stem cell population may be more than just a stochastic phenomenon, but could be a robust strategy [37] that is evolutionary favoured. Correspondingly, understanding the cellular responses requires the consideration of individual cells, from which we obtain information on the diversity of the phenotypic response.

## Results

### Classifying the migratory behaviour of individual HSCs

*Trichinella spiralis* infection provides a model with which to study the variability in responses of individual HSCs. The experimental setup is shown in Fig. 1: HSCs are harvested from healthy control or day 14 infected mice, DiD-labelled, and injected into lethally irradiated recipient mice. *In vivo* imaging is performed approximately 16 hours later for a period of 5 hours. In Fig. 2A-B representative snapshots from the collected movies are shown, together with their corresponding cell tracks given in Fig. 2C.

**Figure 1:**
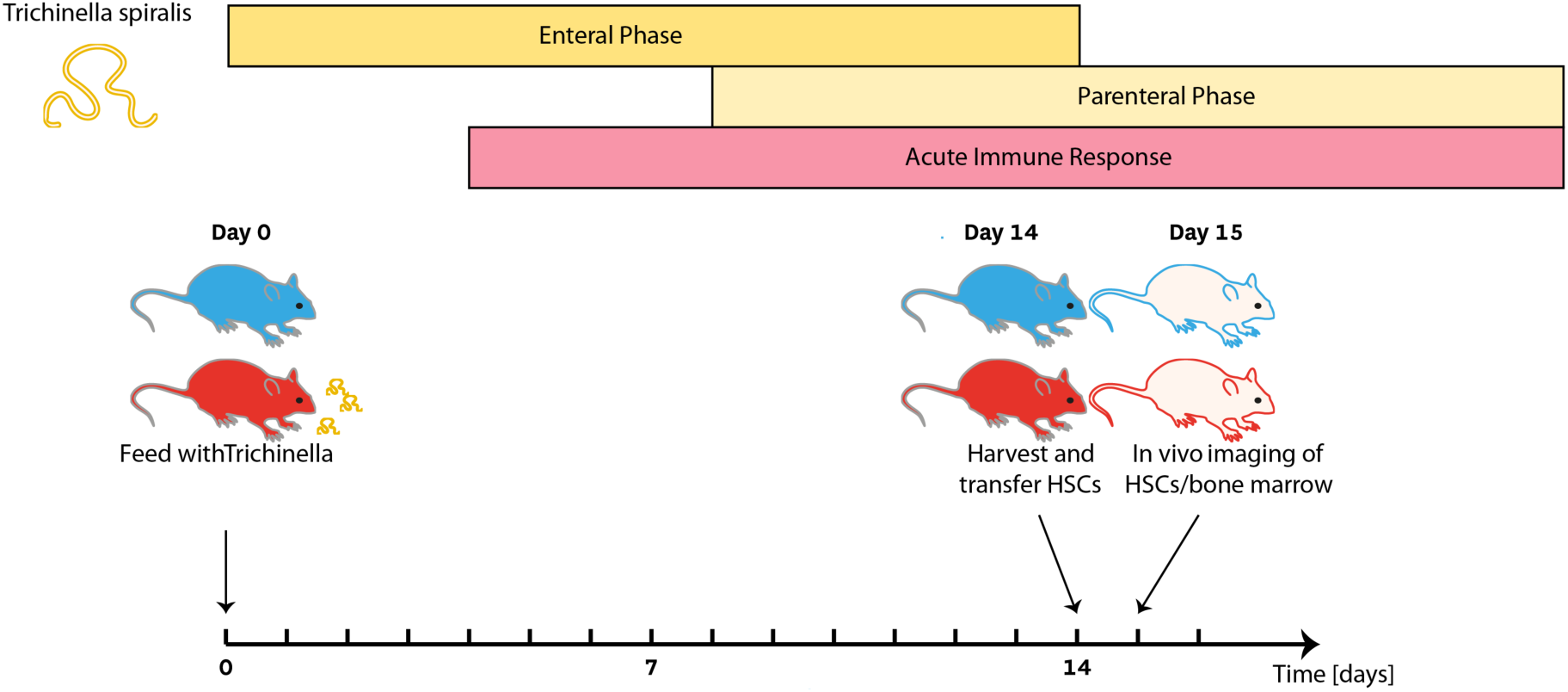
Experimental setup for studying of chronic infection. On day 0 mice are infected by oral gavage (500 larvae each). On day 14 cells are harvested from infected mice and healthy controls and sorted to obtain a population of HSCs. These cells are injected into lethally irradiated healthy recipients, and imaging is performed the following day.

**Figure 2:**
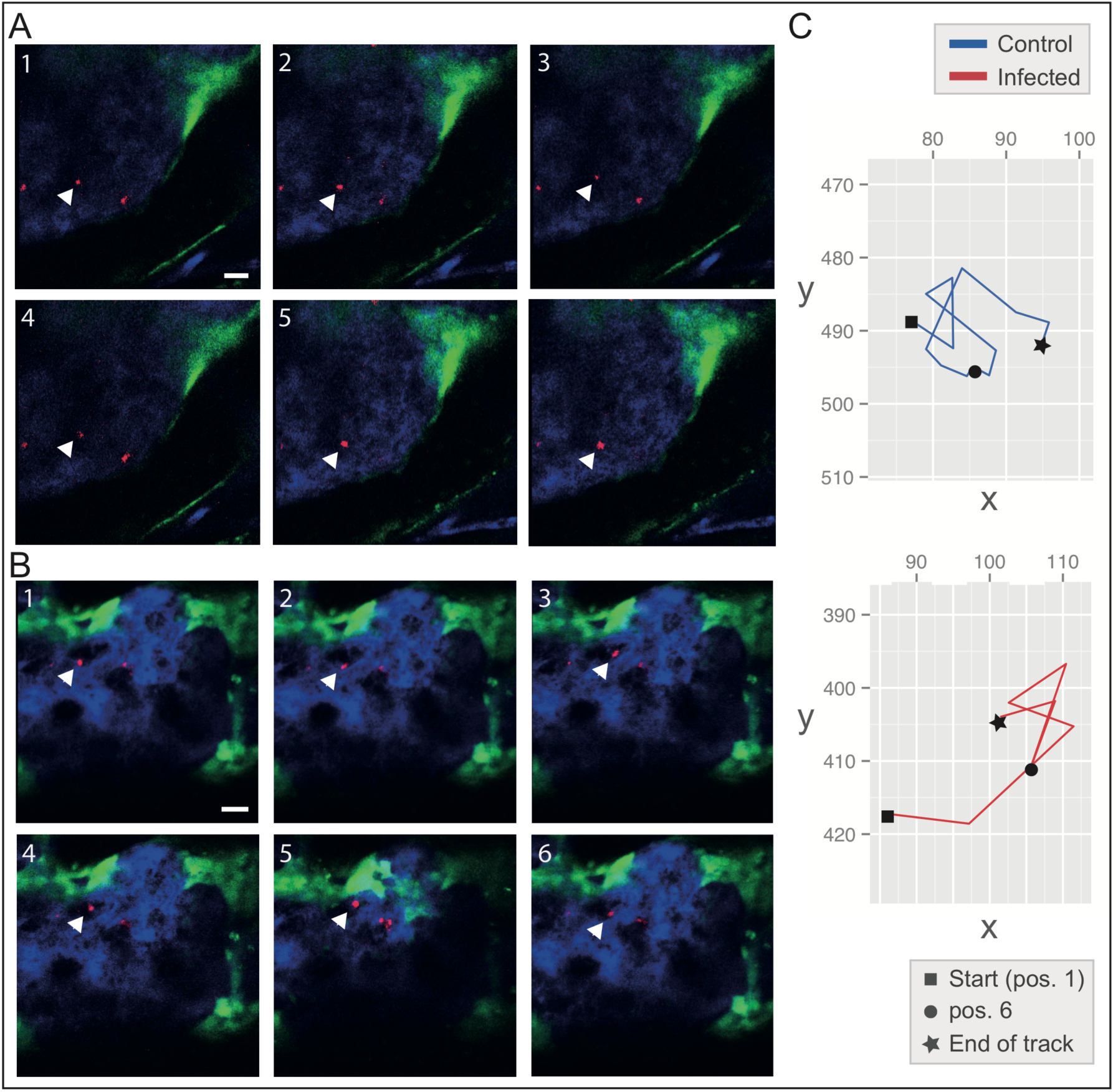
HSC dynamics in the bone marrow microenvironment following Trichinella infection. (A-B) Shown are the first six frames of movies that track HSCs (A) from healthy donor mice and (B) from Trichinella-infected mice. HSCs (red) inhabit a bone marrow niche in proximity to vasculature (blue) and osteoblasts (green). The focal cell being tracked is marked with an arrowhead. Cells were tracked over 5 hours with an interval between frames of 5 minutes; scale bar is 30 μm. (C) Full-length HSC tracks corresponding to movies (A-B) projected from 3D into the x-y plane. Grid positions (in μm) are given relative to the imaging window. In total 26 HSCs from infected mice and 16 HSCs from control mice were analysed.

In order to analyse the behaviour of individual HSCs *in vivo* we reconstruct their 3D migration tracks; rather than using just the centre–of–mass trajectories, we use so-called *α*-shapes (see Methods), which allows us to look at the volumes traversed by HSCs (Fig. S1). We analyse all cells that are found to lie within a total volume of 690 × 580 × 170 *μ*m^3^ inside the bone marrow cavity of the mouse calvarium. The 2D projections of the HSC tracks are shown in Fig. S2. We study the displacement in each dimension in order to quantify the extent of influence of the *Z*-direction on cell behaviour; the variance in this direction is not significantly different from the variance in *x* or *y* (Fig. S2).

In Fig. 3 we show the trajectories (captured by the *α*-shapes) for individual infected and control HSC tracks. The *α*-shape hues reflect the local average curvatures. This enables identification of their convex and concave regions, and thus allows better comparison between the trajectories and the cell behaviours. A convex region indicates that the cell visits its vicinity, partly doubling back on its path; concave curvature, by contrast, suggests forward momentum (or persistence). Here we do find some differences: while we do not find significant differences in concavity (the total proportion of concave regions) between trajectories of HSCs harvested from infected and control mice, we find that *α*-shape surfaces of HSC cells that come from infected mice show fewer transitions between concave and convex regions. This indicates that these surfaces are smoother — either rounder or more elongated — than the *α*-shape surfaces of cells from non-infected mice. This, as may be gleaned from visual inspection of the trajectories in Fig. 3, suggests that the “infected” HSCs either remain local (and very constrained), or exhibit “wanderlust", a pronounced tendency to explore the bone marrow more widely, in search of a different microenvironment. *α*-shapes constructed from infected cells show a significantly different distribution with respect to volumes and surface areas per unit time than those from uninfected cells (Fig. 3 inset).

**Figure 3:**
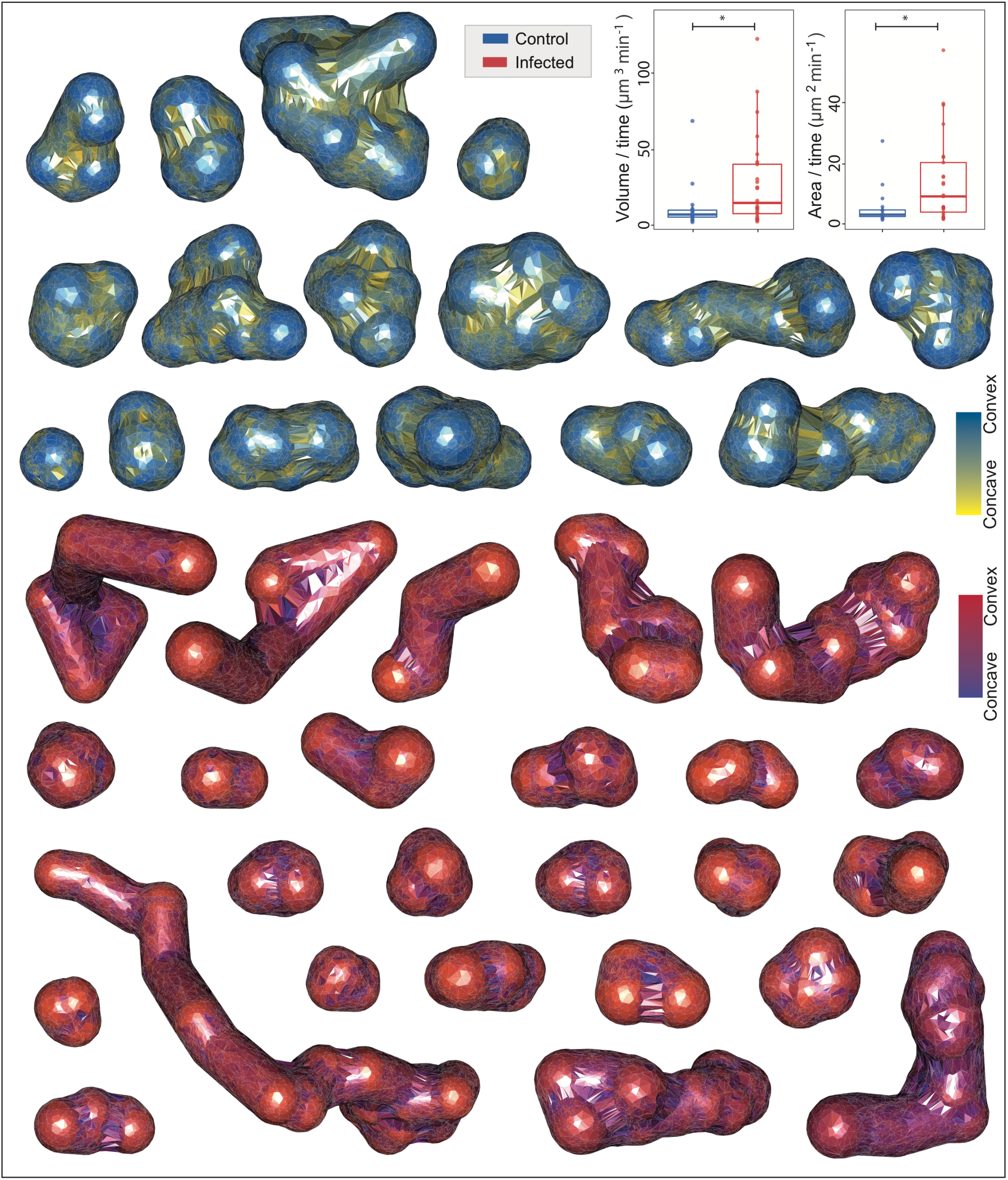
The volumes mapped out by the migration of HSCs via α-shape construction. HSCs originate from either healthy control mice (n = 16, blue) or Trichinella-infected mice (n=26, red). The cells originate from the pooled calvaria of three mice from each category. Infection leads to significant changes in HSC migration behaviour: HSCs are either stationary (or nearly stationary) or become highly migratory. By contrast, HSCs from uninfected mice explore a confined and stable volume within the bone marrow. The inset shows box and dot plots depicting the areas and volumes traversed by the HSCs from infected and control mice. *p < 0.05, according to a Kolmogorov-Smirnov test.

### *T. spiralis* infection leads to heterogenous HSC migration patterns

In order to compare the movement patterns of HSCs from healthy vs *T. spiralis*-infected mice we consider an extensive set of cell migration statistics [23,24]. These are summarised in Fig. 4A: each cell is represented by a point and the population behaviour is additionally represented by violin plots (an extension of the box plot). *Asphericity*, *straightness*, and *sinuosity* each characterise ways in which a track deviates from random (Brownian) motion or from a straight line [38,39] (see Table [tabS1] for full definitions). We compare these statistics between control and infected cells; no significant differences are found. Straightness and sinuosity can both be used to estimate the tortuosity – or “twistedness” – of a path, which can be thought of as the influence of the surroundings on one’s path (for example physical geography on the meanders in a river). The *straightness* of a track (its total displacement/total path length) provides only a cruder estimate of tortuosity (crucially, it only becomes an appropriate statistic for classifying tracks when these get long; so it is not an optimal measure for experiments carried out in animals, where care has to be taken to limit experimental procedures to what is absolutely necessary) since it characterises global straightness and does not account for local differences along the track; *sinuosity* is in general a better estimate of a path’s tortuosity, as it characterises local variation along sections of the track (see [39] for a further details).

**Figure 4:**
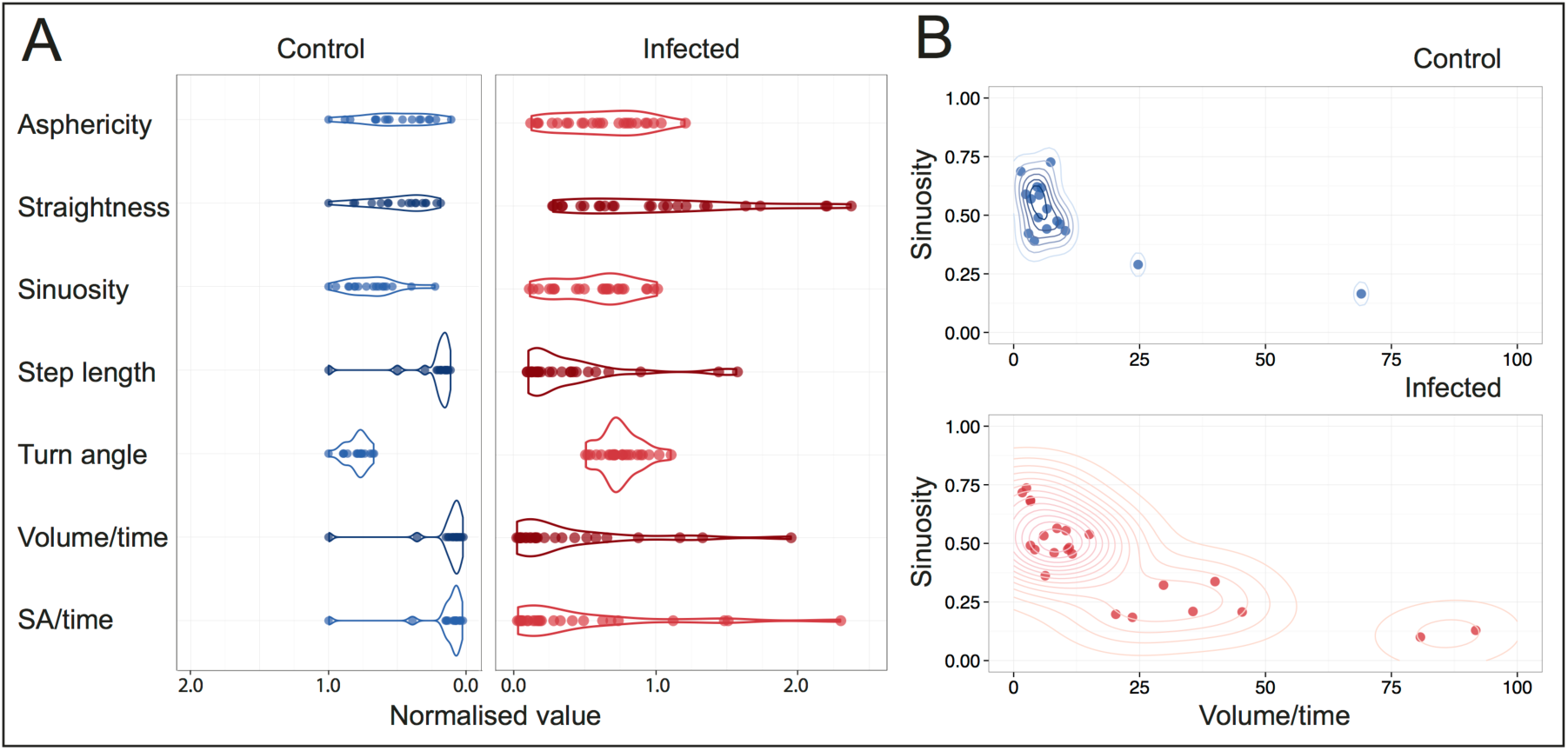
Comparison of cell migration behaviour between control (n=16) and Trichinella-infected (n=26) HSCs. (A) Migratory behaviours are studied via statistics that summarise various aspects of cell movement. For each statistic, the data are scaled by the maximum value from the healthy group; scatterplots with overlaid violin plots of the scatter density are shown. Violin plots are similar to box plots but go further in that they also show the probability density of the data at each point. Step length, volume per unit time, and surface area per unit time are significantly different between the groups (p < 0.05, Kolmogorov-Smirnov test). (B) The joint density plot of two cell-track descriptors that each capture local properties of the track.

The distributions of step length and turn angle show no differences between control and infected cell tracks; but since there is considerable within-track variation for these statistics this is hardly surprising. In Fig 5A we also show the normalised values for the *volume* and *surface area* traversed by each cell per unit time; as reported (Fig. 3), we find statistically significant differences between the distributions of control and infected cells. Here the violin plots help to illustrate both the differences in mean and in variance between the distributions.

In Fig 5B we plot the sinuosity (twistedness) of each cell’s path against the rate at which the cell explores new space (i.e. the volume per unit time). This plot highlights that cells harvested from infected mice show considerably greater variance in each statistic plotted, thus exhibiting a greater range of behaviours than cells from control mice. Comparing the two panels of Fig. 4B suggests that the HSCs collected from infected mice fall into two groups: one, which has the same characteristics as the cells from control mice (high sinuosity and low volume/time), and a second, more dispersed group with lower overall a lower mean value for sinuosity, but a higher mean value for the volume/time. This latter group are much more mobile and migratory in behaviour than the former.

**Figure 5:**
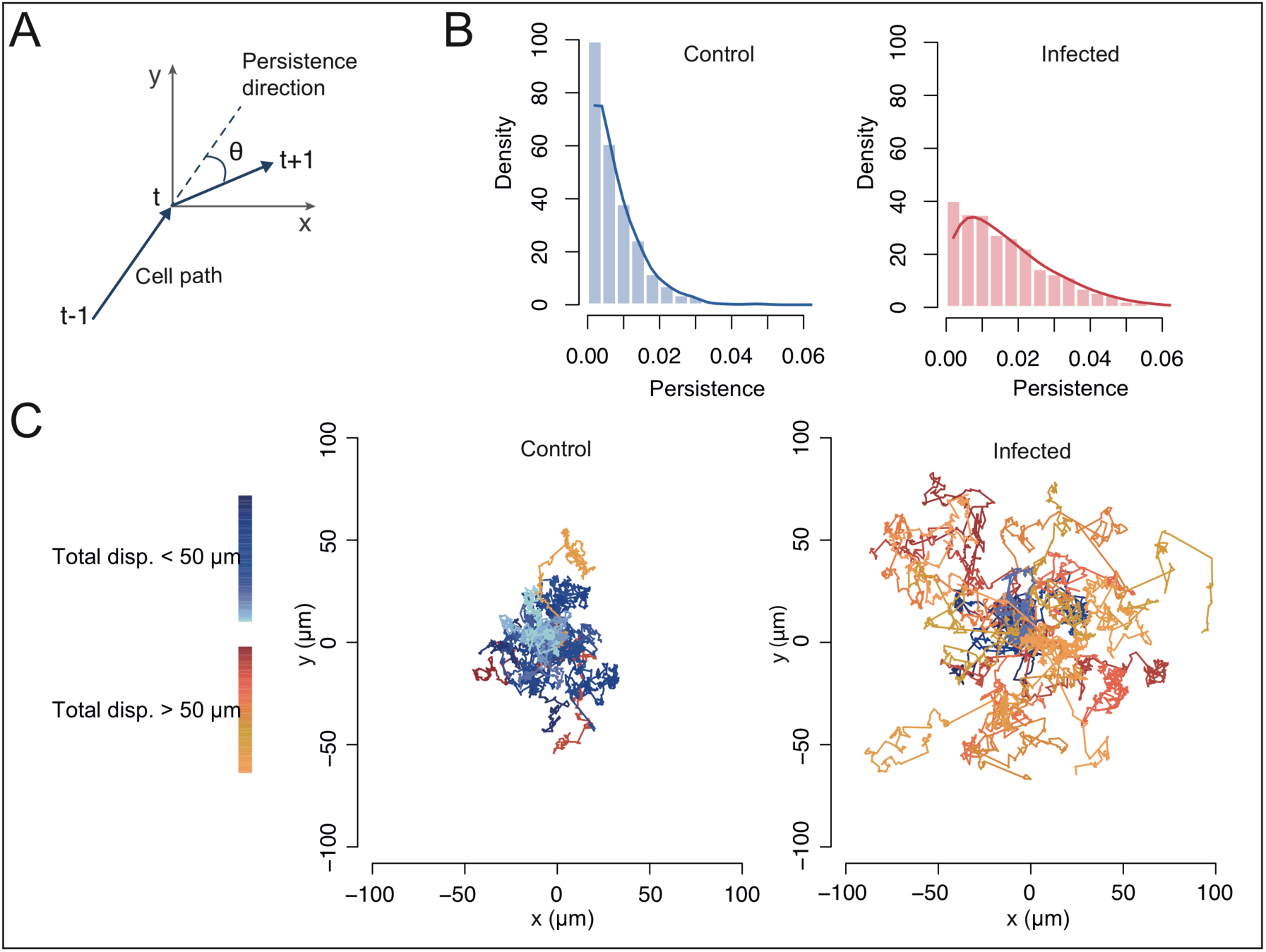
Trichinella-infected HSCs are more persistent than control HSCs and explore greater territories. (A) Schematic description of the random walk model: a walk is simulated in 2D via sampling a step length and a persistence angle θ that quantifies a cell’s deviation from a straight-line path. (B) The inferred level of persistence following statistical inference that combines model and data. (C) Simulations for each group from the inferred model. Each set of simulations contains 20 cell tracks of length 400 steps.

### *T. spiralis* infection leads to more persistent cell movement

Random walk models describe movement (i.e. a path through space) as a sequence of steps and rotations, with, in general, variable step lengths and turning angles. A pure random walk (also known as Brownian motion) is isotropic and displays no overall preference for direction. Other types of random walk are anisotropic by the introduction of bias (a propensity for travelling in a certain direction) or persistence (a propensity for maintaining the direction set by the previous step); models have been developed to describe cell movement in terms of these processes, such as during an immune response [24,40].

A schematic overview of the model we introduce to study persistence, depicting two steps centered at time *t*, is shown in Fig. 5A. We do not study bias here since we cannot (yet) resolve the niche-mediated signals with sufficient spatial resolution to be able to locate them. Thus we assess the contribution of persistence to HSC movement patterns with a random walk model. The level of persistence is inferred by simulation (using Markov Chain Monte Carlo, see details in Methods). Fig. 5B shows the inferred persistence distributions for cells from each group. The level of persistence for each group is small, but the persistence values for HSCs from infected mice are significantly higher overall than those for cells from non-infected mice.

Given the distributions for HSC persistence that we have inferred, we can study the behaviour of HSCs by simulating their movement over time frames that are longer than are experimentally viable to capture. In Fig. 5C we simulate 20 cell tracks for each group by sampling persistence angles from the inferred posterior persistence distributions. We sample step lengths from the empirical distributions of step lengths obtained from tracking data. Whereas we can only track cells *in vivo* for a period of approx. five hours, here we plot the tracks obtained over a simulation period of 33 hours (400 steps), in order to study the longer-term behaviour of these cells. Fig. 5C shows that the relatively small differences in persistence seen between the two groups translates into large cumulative differences in cell migration behaviour over a time frame of around 1.5 days. The distances covered by the cells from the infected group mean that they have the opportunity to encounter, explore, and interact with many more potential niche cells than the non-infected HSCs.

### HSCs perturbed by *T. spiralis* infection display bimodal behavioural patterns

The trajectories shown in Fig. 3 suggest considerable variability in migration behaviour among HSCs collected from *T. spiralis*-infected mice. Particularly, they suggest bimodal behaviour whereby HSCs either occupy confined volumes, perhaps associated with particular niches, with little to no movement; or they become highly migratory and cover large spaces within the bone marrow over the course of the imaging.

In order to investigate this further, we analyse the dynamics exhibited by cells along their trajectories. Again we can make use of the additional information that the *α*-shapes provide and we consider how the cumulative *α*-shapes are generated (Fig. 6A-B). “Domiciliary” cells that are constrained to stay within a limited space (perhaps due to niche supporting factors, or some other, still to be resolved physical limitations) will have a cumulative volume curve with a positive, decreasing gradient that will asymptotically approach 0 because these cells cease to continue to explore new space over time. “Exploratory” cells that are able to move more freely through space, either because of fewer constraints, because of attractant cues, or due to random search behaviour, will have a cumulative volume profile that continues to increase over time.

**Figure 6:**
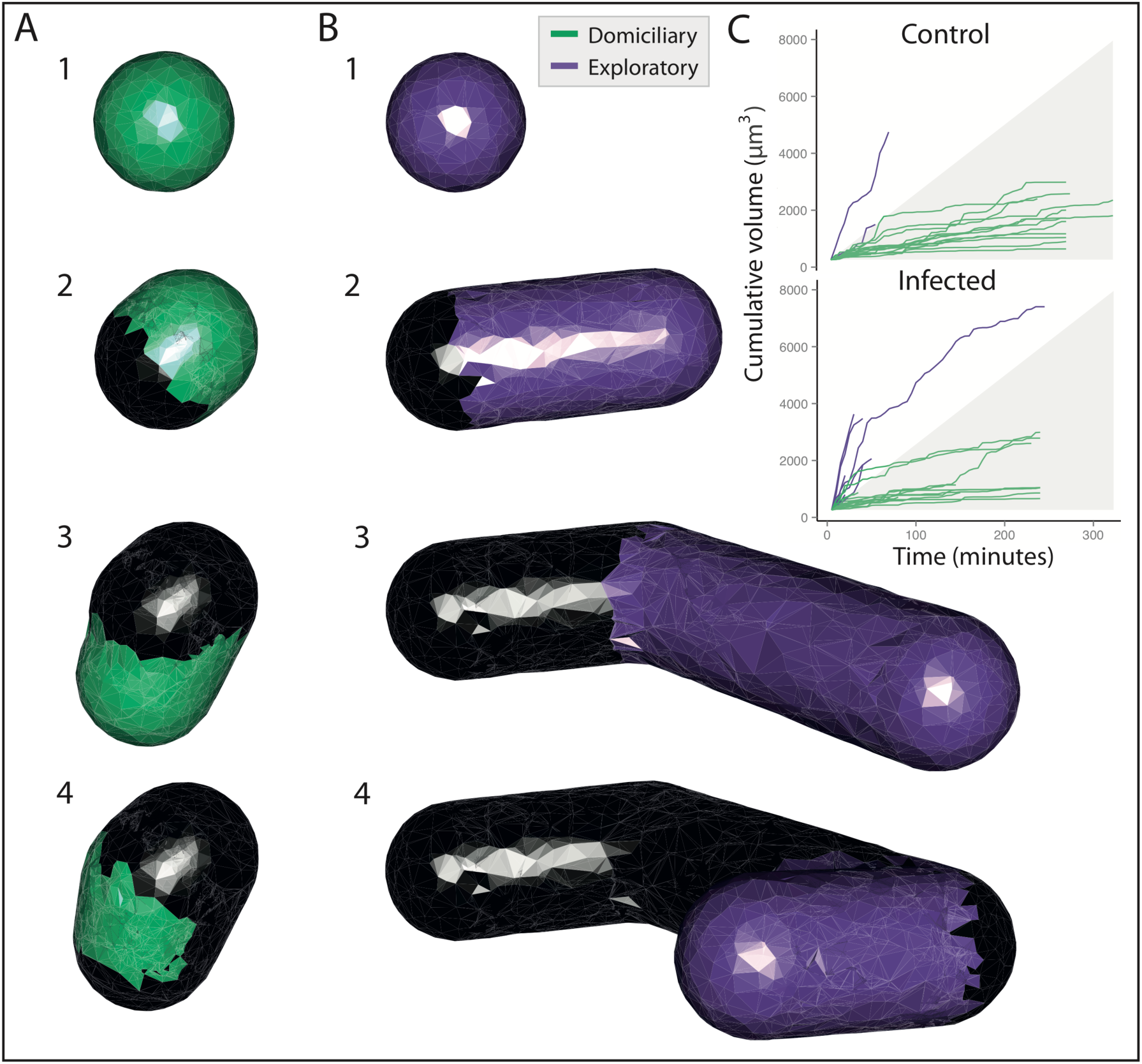
Dynamical evaluation of niche space via α-shapes. (A-B) The first four frames of two α-shapes shows their growth over time. New space visited is shown in colour and space previously occupied is shown in black. (C) Trajectories of α-shape growth depict two growth patterns: domiciliary (green) and exploratory (purple). Shaded region: [0, μdomiciliary + σdomiciliary].

In Fig. 6C the cumulative volume profiles for healthy and infected HSC paths are shown. HSCs from control mice show predominantly the first pattern whereby cells are domiciliary and remain contained within a given volume. In contrast, HSCs from infected mice exhibit two patterns: one group follows the pattern of the healthy cells, maintaining domiciliary tracks, which corroborates what was seen in Fig. 3, where we discern a group of cells that occupy tightly constrained volumes. The second group of cells display markedly different behaviour: these exploratory cells continue to traverse through new space over the course of the tracking period, which leads to the continuing growth of their cumulative volumes.

The relatively homogeneous movement profiles of control HSCs suggest that, at time of imaging (16h post transplantation), they are engaged in stable interactions with a stem cell niche within the bone marrow microenvironment. The heterogeneity of behaviour seen among the cells collected from infected mice suggests some ongoing disruption of the ability to form stable HSC–niche interactions; in other words, infection may have acted to aggravate HSCs, which can no longer engage with niches as successfully. Furthermore, the data suggest that such disruption may occur as follows: in response to the disengagement of HSCs with certain niche factors, the cells adopt either a “cling-on” strategy whereby they do not move away from their current position, or a “abandon ship” strategy that causes them to travel, in search of niche engagement elsewhere.

### The decision of an HSC to migrate from a niche can be understood in terms of its life history

We can draw on life history theory in order to map out when it makes sense for HSCs to become mobile in the bone marrow following an infection (or other perturbation that affects the supportive relationship between a niche microenvironment in the bone marrow and haematopoietic stem cells). Life history theory describes how evolutionary forces shape functional and behavioural aspects of an organism [33,41]. It also provides a framework in which we can study phenotypes from an evolutionary perspective, in order to understand if and how a given behaviour/phenotype may affect an organism’s long-term reproductive potential, i.e. its Darwinian fitness [33]. In addition to canonical uses of life history theory, its ability to describe cellular “life” history choices has been demonstrated in application to stem cells and to cancer [42,43].

Here we use life history theory to study the conditions where it may become beneficial for an HSC to leave its current niche in search for a different niche microenvironment. Benefit is measured as the expected number of long-term (differentiated) offspring cells produced by an HSC adopting a given strategy, or *life-history choice* [33]. Because of the complexity of the niche, its interactions with the HSCs, and the effects of infection on the haematopoietic system, we adopt a general model, that minimises the need for making specific assumptions[44]. We assume that within a niche, *N*_1_ an HSC will produce one offspring at each time step with probability *p*, the probability of division for an HSC. With probability (1 – *p*) the HSC will cease producing offspring (due to loss of stemness or cell death). As we define the fitness of an HSC in terms of its ability to divide, *p* is below referred to as a measure of the fitness of a niche. Shown in Fig. 7A is a schematic of the relative niche fitnesses of two HSC niches, and a possible path between them.

**Figure 7:**
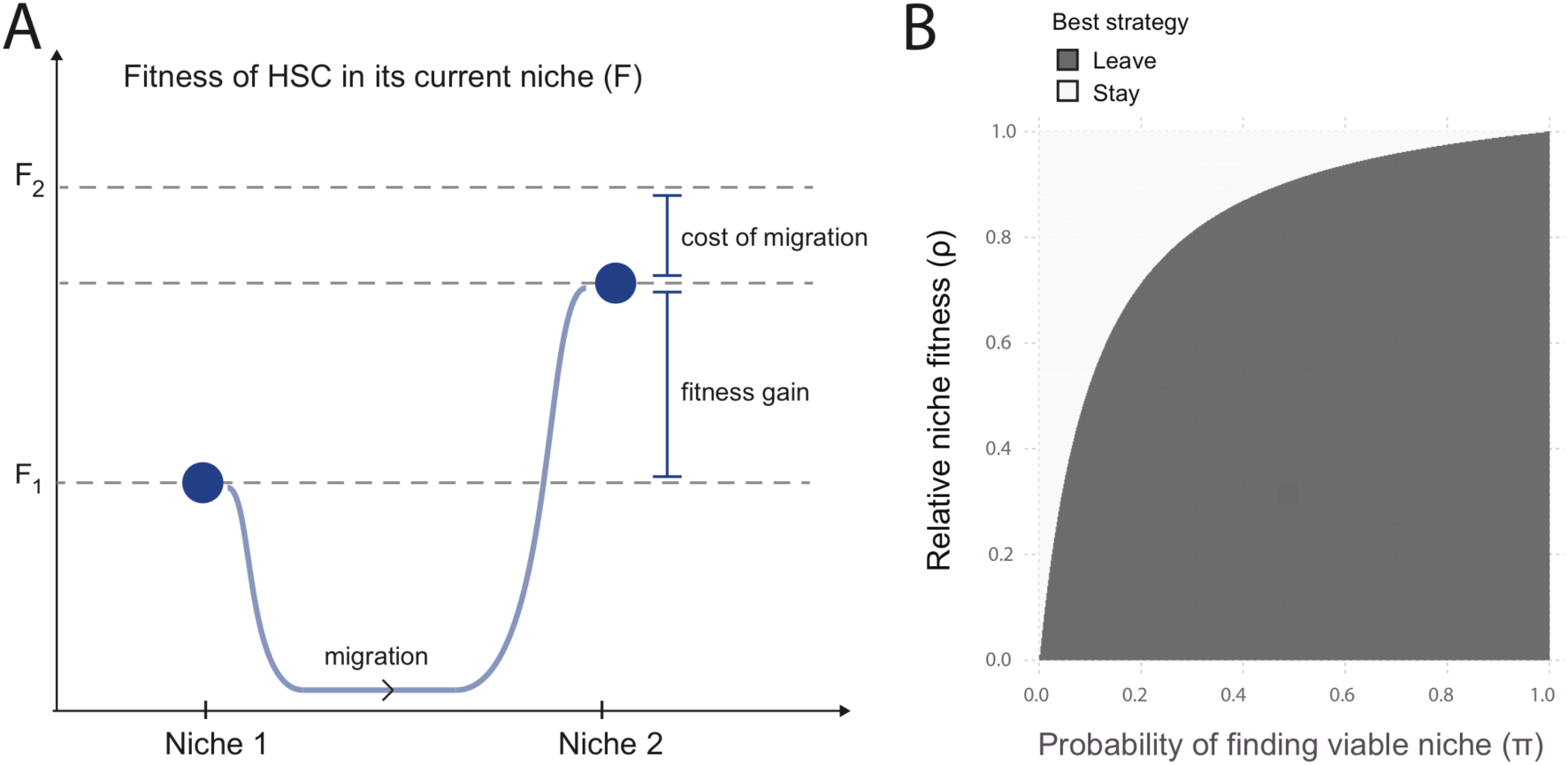
Life history trade-offs describe stem cell dynamics in the bone marrow niche. (A) The blue curve denotes a possible HSC trajectory between two niches N1 and N2: the best strategy is migration, as long as the cost of migration is less than the difference between niche fitnesses. (B) Joint parameter space depicting the best strategy — to stay in or to leave the current niche — under different conditions.

The expected number of progeny, *x*, over the lifetime of the stem cell is then

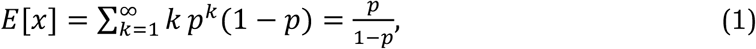

i.e. the number of differentiated cells produced is modelled as a geometric random variable. If the niche deteriorates such that the fitness is *r* = *ρp* with 0 ≤ *ρ* ≤ 1 then we have a concomitant reduction in the expected offspring numbers, *y*, given by

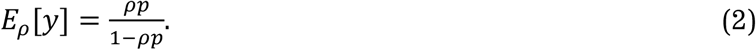

This would be the expected number of offspring in the deteriorating niche were the HSC to stay there. If the probability of finding another functioning niche (with fitness *p*) is *π*, then for

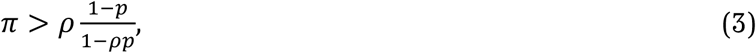

the expected number of offspring for a “wandering” HSC exceeds that of one that stays put in the sub-optimal environment. Here, apart from the assumptions of distinct fitnesses (captured by the fitness ratio *ρ* = *r*/*p*) we have only assumed that a migrating HSC will not seek out other deteriorated niches with fitness *r*, and not move away again from a supportive niche with associated fitness *p*. These assumptions can be relaxed, but this does not, as far as we have tested, alter the qualitative behaviour.

Inspection of Eqn. (3) shows that even for relatively minor deteriorations in the HSC–niche fitness, the *mobile strategy* — i.e. where the HSC becomes migratory — can become advantageous, as long as *π* remains high (as shown in Fig. 7B). Prolonged insult to the bone marrow would, no doubt, lead to a decrease in *π*, i.e. it becomes increasingly less likely to find a niche with associated high fitness, but *ρ* also depends on the spatial location of a niche.

In light of this analysis, the migratory phenotype is likely to be a strategy that can confer considerable advantages to the organism. Maintenance of a healthy haematopoietic system — whether it is in terms of the immune response, or the ability to oxygenate the body — is clearly important to an organism’s reproductive potential. HSCs that react to an infection-driven deterioration in function by going in search of other niches would have, on average, a larger lifetime progeny potential than domiciliary HSCs that do not respond as such. We do not know at this stage whether deterioration in HSC capacity is mediated directly by infection-driven signals, or indirectly, via the niche; however we hypothesise that a niche-driven infection model may be responsible, whereby infection induces changes in the niche that are transduced via niche–stem cell interactions to HSCs. These changes are then maintained upon transplantation and drive the behaviour of transplanted HSCs in new niches, at least transiently.

## Discussion

Analysis of cell behaviour via tracking the migration of cells in 3D is coming of age [23,45]. Significant progress in our understanding of cell morphology, function, and behaviour has been made in recent years due to our ability to study single cells *in vivo* with fluorescent microscopy techniques [46–48]. Here we have provided an in-depth investigation of the 3D migration patterns of HSCs, and discovered that the stem cells exhibit a bimodal response to infection. Haematopoietic stem cells have been subject to intense scrutiny, both due to their essential role in the production of blood, and as a model stem cell system [49,50]. Nonetheless — and at least in parts due to the technical difficulties of imaging HSCs inside the bone marrow — studies that directly visualise HSCs *in vivo* and over time remain rare, despite the insights that they provide [21,22,51].

There is thus little precedent for studying HSC migration and cell movement patterns; this is especially true for the interplay between niche and HSC in cases of physiological stresses on the niche or the haematopoietic system. Previously in [22], an *in vivo* study of stem and progenitor cells, no movement of these cells was observed. Here, elaborating on the results of [21], we show that infection by *Trichinella spiralis* does affect the movement patterns of individual HSCs quite considerably.

Infection affects the haematopoietic system, and HSCs in particular, in both acute and chronic disease. For chronic infections in particular, long-term morbidity due to changes to the cell population of the blood and immune system are amply documented for a range of different infectious diseases. As has been demonstrated [21], the effects of *T. spiralis* infection manifest themselves at the level of HSCs. Here we have analysed this behaviour in more detail and have quantitatively characterised and contrasted the migration patterns of HSCs from infected and uninfected mice.

We base our analysis on the reconstructed trajectories of HSCs (using the *α*-shape approach to describe the 3D trajectory of each imaged HSC) and make use of appropriate statistical descriptors of these cell tracks in order to capture the details of their behaviour. Two central results emerge from this: HSCs collected from control mice show a uniform, largely stationary behaviour. By contrast, HSCs collected from infected mice show what is essentially a bimodal behaviour. Some cells remain stationary and highly localised; the remaining HSCs, however, appear to have entered an “agitated” state, which is reflected by them exploring considerable volumes compared to domiciliary HSCs.

A simple and consistent (but almost certainly not the only possible) explanation is that this behaviour reflects a deterioration of the HSC-niche relationship in the infected mice. In such circumstances, changes to the stem cell may impede its ability to interact with the current niche as is required to maintain its function. Subsequently, an HSC may become agitated enough to move away from the niche. This may either be an active process in order to find an alternative niche environment; or a passive process, if, for example, a niche-retaining signal is lost because of deterioration in the stem cell–niche relationship.

Haematopoietic stem cells must make choices about the migratory behaviour they adopt in a manner similar to the way in which they make choices about division and cell fate [52–54]. It seems natural therefore to consider the migration behaviour of HSCs (and perhaps also myeloid and lymphoid progenitors [55]) as a response to stress in or near to the niche, as well as perhaps a response to a deteriorating niche. Given the characterisation of HSC migration patterns in the bone marrow, a pressing question regards why these stem cells follow the particular movement patterns observed. But our mathematical analysis — building on a large body of work in life history theory — also highlights the distinct possibility that such migration behaviour could be an adaptive, evolutionarily favourable response. As we have shown above for a very parsimonious (as far as possible assumption-free) model of stem cell fate, an HSC that actively leaves a niche in search of a more supportive microenvironment may — under a wide range of scenarios — increase its long-term number of progeny. In other words, if there is a high chance of finding a niche for which a cell is better suited, the proclivity of HSCs to leave their niche should be very high. Most interestingly, and as expected based on our model, we have previously shown that HSCs from *T. spiralis* infected animals have an advantage in reconstituting irradiated recipient mice, as shown by long-term increased engraftment in a competitive bone marrow transplantation setting [21].

In the same vein we can also start to consider the heterogeneity exhibited by HSCs that have been exposed to infection: biological heterogeneity has strong links to robustness. Given the uncertainty of many environments, and the change in demands on an organism these may provoke, employing heterogeneous phenotypes in response appears to be a strategy that has been repeated throughout nature across multiple biological scales [56–58]. Even the existence of certain intermediate progenitor populations has recently been called into question as the role of heterogeneity in the haematopoietic hierarchy grows [59,60]. Our data suggest that the ability of stem cells to respond to a perturbation mediated via infection is founded in part upon the heterogeneity present in the HSC pool within bone marrow.

We can extend these arguments further and explore the predictions that this model makes. These include

- Upon infection we would expect to see depletion of niches that are most easily exposed to a pathogen.
- Some HSCs may even for prolonged infection leave the bone marrow and enter the blood stream.

We would expect that these effects will differ between different infectious diseases, their aetiology, the inflammatory response they elicit, and the way in which their effects manifest themselves among the cells that make up the niche. The effects are also almost certainly not limited to infectious diseases. For example [61] find that periarteriolar niche cells are lost in acute myelogeneous leukaemia-induced sympathetic neuropathy. Or, as another example, Interleukin 27 is an inflammatory marker that has been implicated in changing the size and behaviour of the HSC pool [62]. These lines of evidence are perfectly in line with the first prediction. The second implication of our results is in good agreement with well-established observations that show an increased level of haematopoiesis (also increased numbers of detectable HSCs) in secondary sites of adult haematopoiesis — in particular the spleen — following infection [27]: whereas these sites normally appear to harbour few (if any) HSCs, during infection, HSC activity in the spleen increases considerably, although it is difficult to determine whether these HSCs migrated directly from the bone marrow. We note that the notion of migratory behaviour as an evolutionary favourable strategy has also recently been suggested in ecological settings [63].

This study was able to demonstrate and characterise quantitatively the range of different *in vivo* behaviours of HSCs that have been exposed to an infection, and to contrast this with HSCs collected from control mice. Based on the life history analysis we suggest that the variability in mobility among HSCs collected from infected mice could represent an adaptation and may thus be actively maintained. How inflammatory signals are received and processes by HSCs and/or niche maintaining cells is incompletely understood. Extracting phenotypic data (such as here) and molecular information/profiles (transcriptomic/proteomic) simultaneously is currently not possible. In any case detecting variability at the transcriptional and/or proteomic level between HSCs will require vast numbers of cells, prohibitively so, for *in vivo* experiments in suitable model systems, such as for murine haematopoiesis.

The lack of a meaningful *in vitro* system where such phenomena could be dissected in more detail is both vexing, and — almost certainly — a hallmark of the stem cell’s dependence on its supporting microenvironment. Phenotyping via imaging must be done *in vivo*: the 3D structure of the bone marrow and the complex set of interactions between the niche-maintaining cells/factors crucially affect the behaviour of HSCs, and their response to infection or other perturbations. What may help is an *in silico* niche that allows us to explore how different mechanistic assumptions or hypotheses affect HSC behaviour. Not only would this allow us to place our *in vivo* phenotyping data in the context of available mechanistic knowledge, as well as assumptions, but it would also allow us to make use of modern, simulation-based approaches of experimental design.

## Materials and Methods

### *T. spiralis* infection and in vivo imaging

Lethally irradiated Col2.3GFP mice were injected with approximately 10,000 DiD labelled LT-HSCs (Lin ^*low*^, c-Kit ^+^, Sca1 ^+^, CD34 ^−^, Flk2 ^−^), harvested from the bone marrow of either *T. spiralis*-infected wild type mice or age-matched controls 14 days post-infection (Fig. 1). Sixteen hours post-injection, mouse calvarium were imaged in vivo using multiphoton microscopy, capturing haematopoietic stem cells (HSCs), vasculature and osteoblasts every five minutes for five hours. For full details see [21].

### Cell tracking in 3D

Each imaged calvarium contained markers for HSCs, vasculature and osteoblasts. HSCs were tracked in ImageJ/Fiji [64], using the *TrackMate* plugin, with parameters (estimated diameter, maximum linking distance) set individually for HSCs based on visual inspection of the track, to acquire optimal tracking performance in 3D. HSCs were filtered according to three criteria: roundness, size (< 15 microns in diameter), and composition (single-bodied, i.e. not dispersed DiD objects). Following this filtering process, 26 tracks for infected HSCs and 16 tracks for controls HSCs were retained. Fig. 2 shows snapshot images taken from the movie data for a healthy control (A) or during infection (B), alongside (C) the resulting tracks projected into 2D (for visualisation).

### Construction of *α*-shapes

To represent the niche space occupied by an HSC, we create an *α*-shape modeled on its 3D track [65,66]. The HSC is assumed to have a radius of 4 *μ*m: each 3D coordinate is thus considered to be a sphere of radius 4 *μ*m. Where step lengths were > 4/3*μ*m apart, a cylinder of radius 4 *μ*m was created to connect the spheres. In order to prevent holes in the *α*-shape, inner spheres and cylinders of radius 2 *μ*m were placed within the 4 *μ*m spheres and cylinders. Each inner and outer sphere and cylinder was approximated by a set of 500 points sampled from their surface. The R package *alphashape3d* was used to produce the final *α*-shape objects for each HSC. Fig 3A demonstrates the process of *α*-shape construction for a track composed of two steps. Cumulative *α*-shapes were created by repeating the process of *α*-shape construction sequentially, for each successive time point in the HSC track. In Fig. S1C the convex hull of the track used in Fig. S1A is shown to illustrate the advantage that *α*-shapes have over convex hulls: they represent both the convexity and the concavity of the shape.

### Curvature quantification

To quantify the curvature of an *α*-shape, we adapted methods developed for the analysis of protein surface shape [67]. The *α*-shape vertices connect to form a set of m triangles *T* = {*▯*_1_ *t*_2_, *t*_3_, …, *t*_*m*_}, that constitute the *α*-shape surface. A single vertex *v* is connected to a set of *n* neighbouring vertices in the *α*-shape by the set of connections *C* = {*c*_1_, *c*_2_, *c*_3_, …, *c*_n_}. The curvature *Ω*, calculated for each triangle *τ* in the *▯*-shape, is then given by:

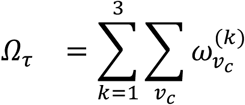

where *ω*_*v*_ denotes the angle formed from two edges joined at vertex *v*, *v*_*c*_ are the set of pairs of connections in *C* such that the resulting triangle is not in *T*, and *k* sums over each vertex in *τ*. This average of angles calculated for each triangle is then used for quantification and visualisation of *α*-shape curvature. A schematic of this procedure is given in Fig. S1B.

### Cell migration statistics

The step length and turn angle distributions are calculated directly from the cell tracks. Volume/time and surface area/time are calculated from the *alpha*-shapes. The straightness index is calculated by *D/L*, where *D* is the maximum displacement and *▯* is the total path length, and gives a global measure of path deviations. The sinuosity index *S* also measures path deviation, but does so around a local region of the path, and provides better estimates of the tortuosity of an undirected random walk than straightness [39]. Sinuosity *S* ∝ σ_*θ*_/*μ*_*x*_, where σ_*θ*_ is the standard deviation of the turn angle distribution and *μ*_f_ is the mean step length. Finally we measure the asphericity of a path by calculating the radius of gyration tensor 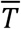 and defining (in 3D) the asphericity *A* as:

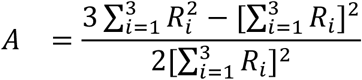

where *R*_*i*_ are the eigenvalues of 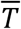.

### Random walk modelling and inference

We use a persistent random walk model to describe stem cell migration [24,40]. The model consists of *N* non-interacting particles migrating in 2D. The direction of a particle’s movement at any time step *t* is described by two random variables, step length *s*_*t*_ and turning angle *θ*_*t*_. A lognormal distribution is used from which to sample step lengths for each particle. The parameters of the lognormal distribution are estimated from the data separately for the control and infected groups.

The turning angle *θ*_*t*_ is defined without loss of generality as the angle between two consecutive motion vectors at time *t*. *θ*_*t*_ follows the wrapped normal distributions [68] with probability density function

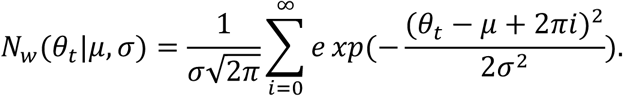

The mean *μ* is 0, while the variance *σ* depends on whether or not the random walker follows persistent motion. The variance for persistent motion is σ_*p*_ = −2*log*(*p*), with persistence parameter *p* ∈ [0,1]. The closer *p* is to 1, the smaller the variances will be, and the more likely the particle will be to sample an angle 0, i.e. continue in the direction of the previous step. If *p* is equal to 0, then the corresponding variance will tend to infinity and the distribution is a wrapped uniform distribution: the cell will exhibit no persistence. At each time point, *θ*_*t*_ and *s*_*t*_ are determined, and the cell moves accordingly a distance of *s*_*t*_ in the direction defined by *θ*_*t*_.

From trajectory data, we infer the parameters of the persistent random walk model in a Bayesian framework for the two treatment groups. The likelihood function (*p*) is derived exactly [24], and we sample the posterior distribution of *p* using a standard Markov Chain Monte Carlo (MCMC) sampler. We run 5000 MCMC steps with the first 3000 discarded as burn-in. We used an adaptive log-Gaussian kernel for each parameter separately with the variance equal to half the variance of all previously sampled parameter values.

In order to determine HSC behaviour over longer time intervals than is experimentally feasible or justifiable (see Experimental design in the next section), we simulate 50 trajectories based on the inferred posterior distribution of the persistence and the estimated step length distributions for each group. All simulations were run for 400 time points, corresponding to 33.3 h, starting from the origin.

### Experimental design and analysis

Guidelines demand that we minimise the number of experiments in animals to a level where statements of statistical significance can be made. Often a better analysis and improved experimental design can vastly increase our power to discern different types of behaviour (at the same effect size), as we demonstrate here.

For the analysis of cell migration behaviour, we can use this to both reduce the number of animals studied, and refine the procedure by developing better statistical analyses that allow us to identify patterns reliably in a shorter imaging window. Analysis of cell migration data is naturally simpler the longer we can track cells; different modes of cell migratory behaviour can be discerned much more readily from data that contain a large number of long cell trajectories. Here, in order to minimise observation time, we have focussed on those aspects of cell tracks that allow us to discriminate between different types of behaviour but that do not require a global characterisation of the paths of the cells (see Table [tabS1]). For these measures the collected data are sufficient to contrast the behaviour of cell migration of HSCs taken from control and infected mice, and to highlight statistically significant differences between the two groups.

In addition, we performed calculations of statistical power in order to estimate the sample sizes that would be required in order to observe statistical significance for the effects seen in other migratory statistics analysed. For the step length distribution, approximately 59 cells of each group would be required to show that the mean is different between control and infected cells, and for the turn angle the equivalent group sample size is 98. We note that we are testing the null hypothesis of equal means – rather than equal distributions here – and also we are assuming that the variance is known and equal between groups, an assumption which, given the heterogeneity that we have observed so far, we think may well not hold in practice.

## Acknowledgments

The authors would like to thank Marc Mangel for comments on the manuscript. Funding is gratefully acknowledged from the Biotechnology and Biological Sciences Research Council, BB/L023776/1 (ALM, RK, CLC, MPHS), and the National Centre for the Replacement, Refinement and Reduction of Animals in Research, NC/K001949/1 (JL). The funders had no role in study design, data collection and analysis, decision to publish, or preparation of the manuscript.

## Competing Interests

The authors declare that no competing interests exist.

---

**Figure S1:**
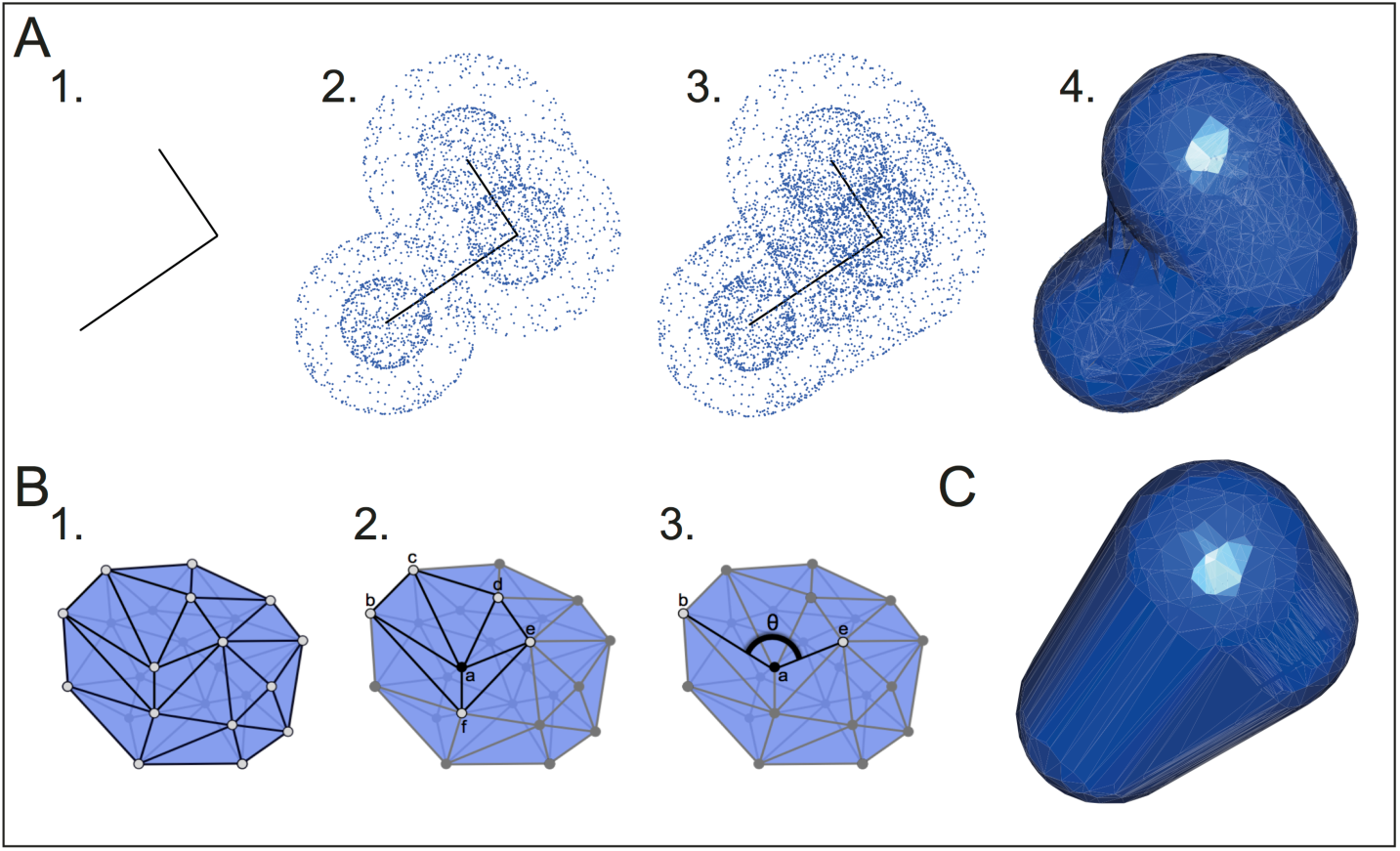
A model of 3D niche space. (A) (1.) An initial track (of two steps in length) provides a skeleton around which (2.) points are sampled on spheres of radius 4μm and 2μm. (3.) Cylindrical sampling is used to connect spheres with separation > 4 μm. (4.) Rendering of the resultant α-shape. (B) Curvature is quantified by the following procedure: (1.) for each vertex (grey dot) in the α-shape, (2.) find its neighbours, and (3.) sum the angles formed by each pair of connections, for all sets of connections that do not form a triangle on the surface of the shape. (C) The corresponding convex hull for the trajectory shown in (A).

**Figure S2:**
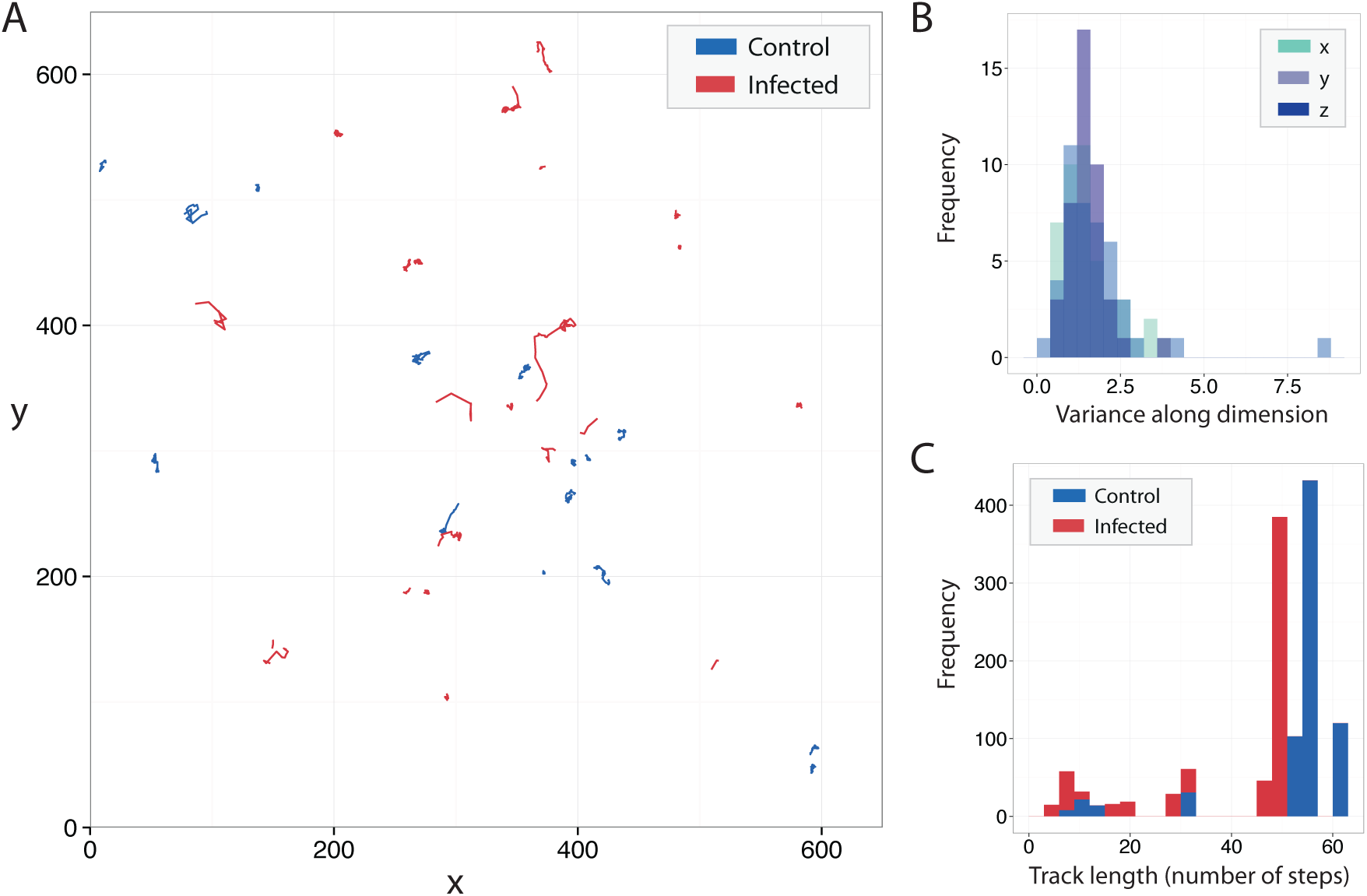
Characteristics of cell tracks. (A) Projection of 3D tracks into the xy-domain. (B) Comparison of the variance in each dimension demonstrates that the z-direction is not significantly different from x or y. (C) Histogram of the cell track lengths for control and infected groups.

**Table S1.** Stem cell migration statistics for ctl. tracks. Description of the cell migration statistics used to quantify tracks in 3D for control (ctl.) cells. SA, surface area.

| Statistic | Description | Ctl. Range | Ctl. Mean |
| --- | --- | --- | --- |
| Asphericity | 'Directedness' of the path | [0.0891, 0.810] | 0.420 |
| Straightness | (Total distance)/(total path length) | [0.0694, 0.374] | 0.185 |
| Sinuosity | 'Twistedness' of the path | [0.165, 0.726] | 0.50 |
| Step length | Distance between consecutive steps | [1.04, 9.12] | 2.20 |
| Turn angle | Angle between consecutive motion vectors | [1.53, 2.28] | 1.82 |
| Volume/time | Volume occupies per unit time | [1.42, 69.0] | 10.4 |
| SA/time | SA occupies per unit time | [0.664, 26.7] | 4.23 |

**Table S2.**
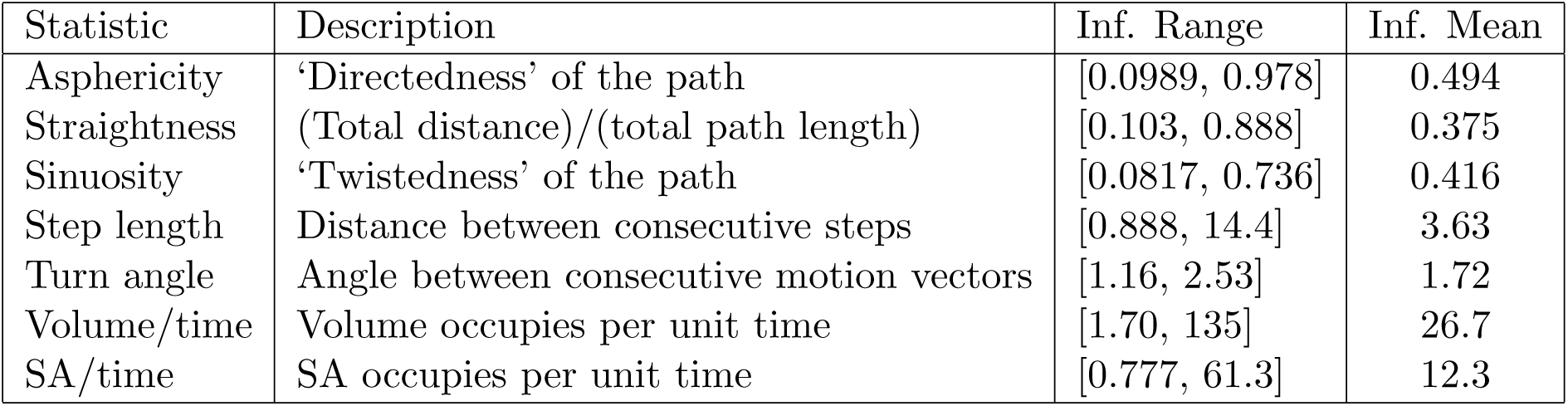
Stem cell migration statistics for inf. tracks. Description of the cell migration statistics used to quantify tracks in 3D for infected (inf.) cells. SA, surface area.

